# Extensive farming in Estonia started through a sex-biased migration from the Steppe

**DOI:** 10.1101/112714

**Authors:** Lehti Saag, Liivi Varul, Christiana Lyn Scheib, Jesper Stenderup, Morten E. Allentoft, Lauri Saag, Luca Pagani, Maere Reidla, Kristiina Tambets, Ene Metspalu, Aivar Kriiska, Eske Willerslev, Toomas Kivisild, Mait Metspalu

## Abstract

Farming-based economies appear relatively late in Northeast Europe and the extent to which they involve genetic ancestry change is still poorly understood. Here we present the analyses of low coverage whole genome sequence data from five hunter-gatherers and five farmers of Estonia dated to 4,500 to 6,300 years before present. We find evidence of significant differences between the two groups in the composition of autosomal as well as mtDNA, X and Y chromosome ancestries. We find that Estonian hunter-gatherers of Comb Ceramic Culture are closest to Eastern hunter-gatherers. The Estonian first farmers of Corded Ware Culture show high similarity in their autosomes with Steppe Belt Late Neolithic/Bronze Age individuals, Caucasus hunter-gatherers and Iranian farmers while their X chromosomes are most closely related with the European Early Farmers of Anatolian descent. These findings suggest that the shift to intensive cultivation and animal husbandry in Estonia was triggered by the arrival of new people with predominantly Steppe ancestry, but whose ancestors had undergone sex-specific admixture with early farmers with Anatolian ancestry.

## Introduction

The change from hunting and gathering to farming was associated with important demographic and cultural changes in different parts of the world. The process involving changes in life-style and in material culture, such as the introduction of pottery, has often been referred to as the Neolithic transition. This term can become confusing when talking about Eastern and Northern areas of Europe like Estonia where some aspects of the so-called Neolithic package, such as pottery, arrive earlier, whilst the transition from hunting-gathering to farming as the main source of sustenance does not occur until Late Neolithic^1^.

aDNA studies have shown that Mesolithic hunter-gatherer groups of Europe and West Asia were genetically highly differentiated from each other in terms of genetic distances estimated from autosomal loci^2–4^. At the same time, and likely as a result of multiple population turnovers and bottlenecks^5^, the Mesolithic hunter-gatherers of Europe displayed a highly uniform pallet of mtDNA with most individuals belonging to haplogroup (hg) U5^6,7^. The early farmers of Europe, who arrived from West Asia more than 8,000 years BP (yr BP)^8,9^, in contrast belonged to a wide variety of mtDNA haplogroups previously unseen in Europe^10,11^. These findings have led to the view that the shift to agriculture involved a substantial degree of population replacement and that hunter–gatherers and farmers did not interbreed considerably for the first few thousand years, both in Central Europe^12–14^ and in Scandinavia^15–17^. Analyses of autosomal data on a genome-wide scale have revealed that among the present-day populations of Europe, Sardinians show the highest affinity to the early farmers of Europe. The ancient genomes of the earliest farmers from across Europe – including those from Scandinavia^15,16^, the Alps^18^, Central^2,19,20^ and Southern Europe^2,21^ – cluster closely together and share genetic components with present-day Sardinians and ancient DNA sequences from Anatolian farmers suggesting their common descent from an Anatolian source^4,22^. It has also been shown that the first farmers of the Fertile Crescent had a clear regional substructure largely copying that of the local hunter-gatherers^4^. Importantly, the early farmers of Europe derive from Anatolian farmers^4,23^ and not from the eastern side of the Fertile Crescent^3,4^. The Yamnaya Culture people from the Steppe region of the Eastern European Plain associated with the next wave of migration into Central Europe^2,24,25^ on the other hand shared some ancestry with hunter-gatherers of Caucasus and Iran^4^.

The first signs of crop cultivation in Estonia appeared 6,000 yr BP, 2000 years after the first evidence of farming in Southern Europe, while the transition to the intensive cultivation and animal husbandry is estimated to have taken place much later, between 4,800–4,000 yr BP^26^. This change is associated with a gradual shift from the primarily hunting-gathering-based Comb Ceramic Culture (CCC) to the farming-based Corded Ware Culture (CWC). CCC is thought to have arisen around 5,900 yr BP in areas east of the Baltic Sea and spread as far as Estonia, parts of Finland, Sweden, Russia, Belorussia, and Latvia and Lithuania (Fig 1)^27^. CCC sites have been found predominantly near bodies of water and it has been suggested that the hunter-gatherer groups, making clay pots that were decorated with a comb-like stamp, largely lived off of fishing, hunting and gathering, as demonstrated by the animal bones found from CCC sites and by stable isotopes (^13^C and ^15^N) of human remains^27–29^. Conversely, the rise of CWC around 4,800 yr BP^30^ has been associated with the late spread of farming. CWC later reached Finland, Sweden and Norway in the North (Fig 1), Tatarstan in the East, Switzerland and Ukraine in the South, and Belgium and France in the West^31–33^.Demic diffusion from the Steppe region^34^ has been recently supported by aDNA evidence and proposed to be associated with the appearance of CWC in Europe^2,25^. CWC in Estonia is, among other traits, characterized by clay vessels often decorated with cord impressions, boat-shaped stone axes^27^ and clear-evidence of cultivation and animal husbandry, i.e. bones of sheep/goat, pig, and cattle as well as artefacts made of them found mostly from burial sites, numerous occurrences of *Cerealia* pollen in bog and lake sediments, barley seed and a seed imprint on a pot shard, and stable isotopes (^13^C and ^15^N) of human bones^35–37^. Genetic studies from Central Europe have revealed that CWC burials reflected their familial bonds^38^ and that these Late Neolithic farmers had already admixed with hunter-gatherers, and also brought novel mtDNA haplogroups to the region^12^. They were found to be most similar to modern populations from Eastern and Northern Europe, and the Caucasus^12,25^. Despite its pivotal role in the Yamnaya/CWC expansion, the Eastern European region has been largely overlooked in ancient DNA studies until very recently^39^. Furthermore, as a consequence, there is very little information about the genetic processes involved in farming reaching this area. Additionally, of all modern European populations analysed so far, East Europeans and particularly Estonians are the ones with the smallest fraction of the Anatolian Neolithic genetic component^2^, pointing to a potentially different process of farming reaching this area.

**Fig 1.**
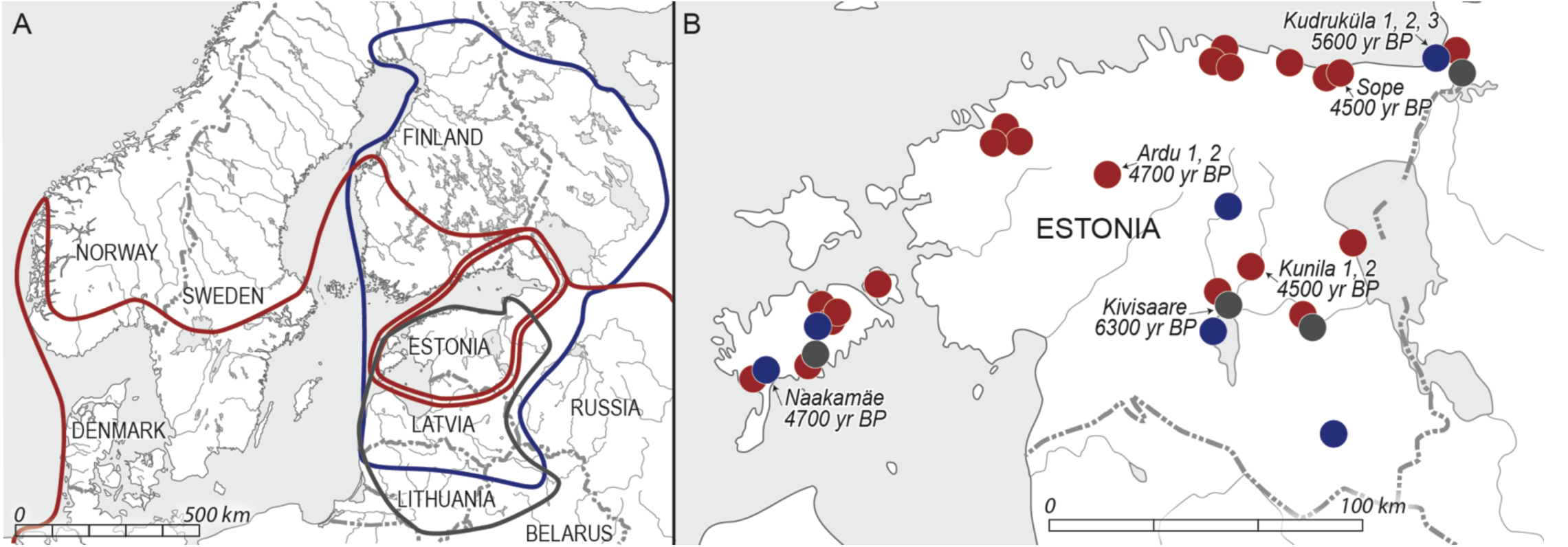
Map of the spread (A) and burial sites (B) of the archaeological cultures involved in this study. Narva Culture is marked with grey, Comb Ceramic Culture with dark blue and Corded Ware Culture (CWC) with dark red. A. Double red line indicates an archaeologically distinct CWC area. B. Sites involved in this study are indicated with arrows along with numbers of analysed samples and the approximate mean values of calibrated carbon dating results.

To shed more light on the genetic changes during the shift to farming based economies in Estonia, we extracted and sequenced aDNA from skeletal remains uncovered in the context of Mesolithic Narva Culture (NC) (7,200–5,900 yr BP), and Neolithic CCC (5,900–3,800 yr BP) and CWC (4,800–4,000 yr BP) from Estonia (Fig 1). We compared these data to sequence and genotype data from modern and ancient populations of Europe, West Asia and Siberia to make inferences about the extent of continuity and genetic change during the end of the Stone Age in Estonia.

## Results and Discussion

DNA was extracted from teeth from skeletal remains of ten individuals from Estonia dating to 4,500–6,300 yr BP. These included five individuals associated with the Neolithic CWC, four with the Neolithic CCC and one with the Mesolithic NC (Fig 1; Table 3). Initial shotgun sequencing yielded endogenous DNA (reads mapping to the human genome) in proportions higher than 1% for eight individuals whose genomes were further shotgun sequenced to whole genome coverage between 0.01 and 2.13 (Table 4). Genetic sexing based on the proportion of reads mapping to X chromosomes confirmed assignments based on morphology (available for 6/10 samples) and provided additional information about the sex of the individuals involved in the study (Table 4).

**Table 3.**
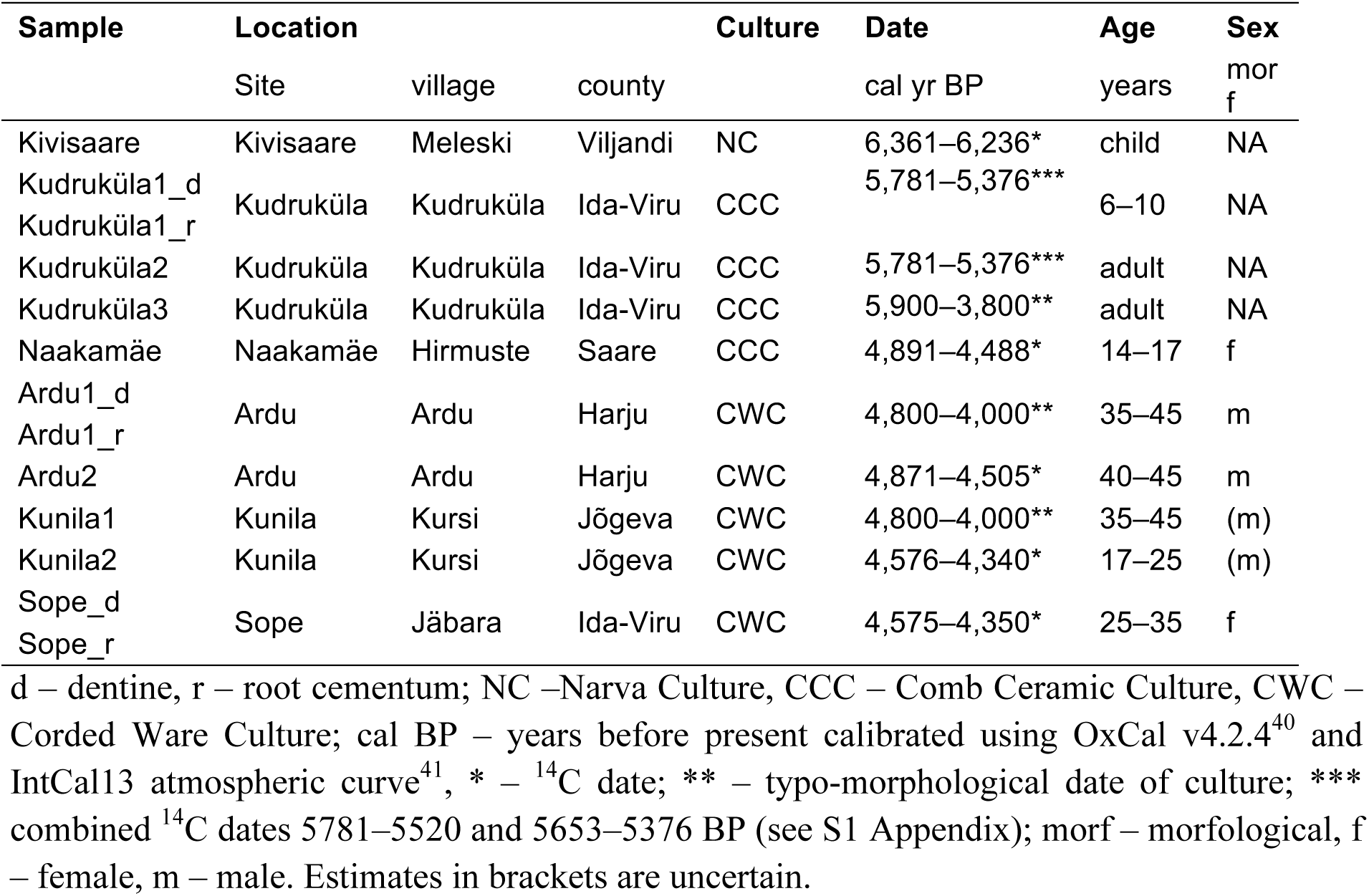
Sample information.

**Table 4:** Sequencing results, genetic sex, DNA damage and contamination estimates of the samples of this study.

| Sample | Culture | Sex<br>morf | Sex<br>gen | Endo<br>% | Cov<br>x | MT<br>cov<br>x | C=>T<br>% | $\lambda$<br>% | $\delta s$<br>% | MT cont<br>% [CIs] | Male X cont<br>% |
| --- | --- | --- | --- | --- | --- | --- | --- | --- | --- | --- | --- |
| Kivisaare | NC | NA | XY | 1.94* | 0.01 | 5.66 | 23.2 | 37.7 | 64.0 | 1.6 [37.1;0.3] | NA |
| Kudruküla1_d | CCC | NA | (XX) | 0.24 | 0.0006 | 6.73 | 10.7 | 34.9 | 30.0 | 0.7 [13.3;0.1] | – |
| Kudruküla1_r |  |  | (XX) | 0.60 | 0.0006 | 1.41 | 12.5 | 38.7 | 35.5 | NA | – |
| Kudruküla2 | CCC | NA | XX | 7.02* | 0.07 | 10.39 | 8.3 | 36.0 | 22.8 | 0.3 [6.9;0.1] | – |
| Kudruküla3 | CCC | NA | XY | 2.27* | 0.11 | 14.29 | 14.1 | 34.3 | 37.6 | 0.3 [3.3;0.0] | NA |
| Naakamäe | CCC | f | NA | 0.02 | 0.00006 | 0.07 | 12.5 | 39.0 | 31.9 | NA | – |
| Ardu1_d | CWC | m | XY | 34.20* | 0.25 | 77.08 | 15.8 | 31.0 | 37.1 | 1.3 [2.4;0.5] | (0.6/2.6) |
| Ardu1_r |  |  | XY | 79.98 | 0.50 | 98.72 | 7.5 | 31.0 | 17.6 | 0.7 [1.5;0.2] | NA |
| Ardu2 | CWC | m | XY | 82.00* | 1.02 | 70.31 | 10.3 | 44.7 | 31.0 | 1.3 [2.5;0.5] | 1.1/0.9 |
| Kunila1 | CWC | (m) | XY | 30.07* | 0.58 | 58.67 | 23.7 | 38.3 | 66.1 | 2.5 [4.4;1.4] | 1.4/1.9 |
| Kunila2 | CWC | (m) | XY | 60.63* | 2.13 | 129.52 | 24.1 | 38.0 | 65.5 | 0.2 [0.9;0.0] | 1.3/0.9 |
| Sope_d | CWC | f | (XX) | 0.31 | 0.0002 | 1.24 | 15.5 | 38.3 | 42.4 | NA | – |
| Sope_r |  |  | XX | 11.70* | 0.30 | 90.48 | 14.6 | 44.0 | 46.1 | 0.1 [1.2;0.0] | – |

d – dentine, r – root cementum; NC –Narva Culture, CCC – Comb Ceramic Culture, CWC – Corded Ware Culture; morf – morfological, f – female, m – male; gen – genetic; Endo – endogenous DNA, \* –sequenced further; Cov – genomic coverage; MT cov – mitochondrial DNA coverage; MT cont – mitochondrial DNA contamination; Male X cont – X chromosome contamination for males. Estimates in brackets are uncertain.

### mtDNA diversity of Estonian Stone Age hunter-gatherers and farmers

Mitochondrial DNA coverage was sufficiently high for haplogroup assignments for 9 out of the 10 individuals (S1 Table). The only individual that had to be excluded from further analyses because of low endogenous DNA content and coverage was one CCC individual from the Naakamäe site. The NC individual and all three successfully haplogrouped CCC individuals belonged to hg U (more specifically sublineages of U5a, U5b, U4a and U2e) (Table 1). The CWC individuals displayed a more diverse set of mitochondrial haplogroups, including H5a, T2a and J1c (Table 1). Notably 2 out of 5 CWC individuals shared hg U5b affiliation with the hunter-gatherer groups (Table 1).

**Table 1.** mtDNA haplogroups of the studied samples.

| Sample | Culture | Date (cal yr BP) | MT hapl |
| --- | --- | --- | --- |
| Kivisaare | NC | 6,361–6,236* | U5a2d |
| Kudruküla1_d | CCC | 5,781–5,376*** | U5b1d1 |
| Kudruküla1_r |  |  | U5b1d1 |
| Kudruküla2 | CCC | 5,781–5,376*** | U4a |
| Kudruküla3 | CCC | 5,900–3,800** | U2e1 |
| Naakamäe | CCC | 4,891–4,488* | NA |
| Ardu1_d | CWC | 4,800–4,000** | T2a1a |
| Ardu1_r |  |  | T2a1 |
| Ardu2 | CWC | 4,871–4,505* | U5b2c |
| Kunila1 | CWC | 4,800–4,000** | U5b1b |
| Kunila2 | CWC | 4,576–4,340* | J1c3 |
| Sope_d | CWC | 4,575–4,350* | H5a |
| Sope_r |  |  | H5a1 |

d – dentine, r – root cementum; NC – Narva Culture, CCC – Comb Ceramic Culture, CWC – Corded Ware Culture; cal BP – years before present calibrated using OxCal v4.2.4<sup>40</sup> and IntCal13 atmospheric curve<sup>41</sup>; \* – <sup>14</sup>C date; \*\* – typo-chronological date of culture; \*\*\* – combined <sup>14</sup>C dates 5781–5520 and 5653–5376 BP (see S1 Appendix); MT hapl – mitochondrial DNA haplogroup.

The prevalence of hg U among Estonian CCC individuals is in accordance with previous results on European hunter-gatherer populations^5–7,12,13,15–17^. Thus Estonian CCC was a population that likely had not yet had significant sexual contact with the early farming populations of Europe. In contrast, three of the CWC individuals from Estonia belonged to haplogroups that first appeared in Europe during the Neolithic (H5, J1 and T2)^10–17^. Thus, while the sample size in this study is arguably small for drawing firm conclusions based on one non-recombining locus, our results do suggest that the transition to farming and animal husbandry in Estonia, which occurs with the arrival of CWC, was mediated by immigration of people.

The nine Estonian ancient mtDNA lineages were compared against a dataset of 2291 modern Estonian mtDNA sequences and found to map within the modern Estonian mtDNA variation (number of differences from the closest modern sample 0–9, mean 3.2) (S1 Figure; S2 Figure).

### Y chromosome uniformity of Estonian Corded Ware samples

Y chromosome haplogroup affiliation could be determined for 5 male individuals with whole genome coverage of 0.1x or above (Table 2). While the chances that a position of interest is sequenced at this low genome-wide coverage are marginal, the basic haplogroup assignments are still robust due to the multitude of equivalent markers that define each clade. Based on the available markers (Table 2) all five individuals could be confidently assigned to hg R and none to hg N, which is a highly common haplogroup in modern Estonians (31%)^42^.

**Table 2.** Y chromosome haplogroup affiliations of Estonian Corded Ware Culture (CWC) farmers and a Comb Ceramic Culture (CCC) hunter-gatherer.

| Sample | Culture | Haplogroup-<br>marker | N<br>M231 | R1<br>M173 | R1a<br>M420 | R1a<br>M459 | R1a5<br>YP1272 | R1a<br>M198 | R1a<br>M417 | R1a<br>Z645 | R1a1<br>Z283 |
| --- | --- | --- | --- | --- | --- | --- | --- | --- | --- | --- | --- |
| Kudruküla3 | CCC | R1a5-YP1272 | 12\|0 | 1\|7 | 0\|2 | 0\|0 | 2\|1 | 4\|0 | 2\|0 | 0\|0 | 0\|0 |
| Ardu1_r | CWC | R1a-Z645 | 45\|1 | 0\|44 | 0\|10 | 0\|3 | 6\|0 | 0\|10 | 0\|4 | 0\|2 | 0\|0 |
| Ardu1_d | CWC | R1a-Z645 | 32\|0 | 1\|21 | 1\|1 | 0\|0 | 5\|0 | 0\|4 | 0\|2 | 0\|1 | 0\|0 |
| Ardu2 | CWC | R1a-Z645 | 68\|0 | 2\|58 | 1\|9 | 0\|7 | 17\|0 | 0\|7 | 1\|7 | 0\|1 | 0\|0 |
| Kunila1 | CWC | R1a-Z645 | 53\|1 | 0\|45 | 0\|8 | 0\|6 | 9\|0 | 1\|11 | 0\|4 | 0\|2 | 0\|0 |
| Kunila2 | CWC | R1a1-Z283 | 115\|1 | 2\|105 | 4\|18 | 2\|12 | 23\|0 | 2\|12 | 3\|7 | 0\|2 | 0\|1 |

The number of haplogroup defining sites^43^ covered by at least one read in our data, is shown on the branches of the tree drawn above the table. In each cell of the table the number of sites detected in the given sample in ancestral (a) versus derived (d) state is shown in the format 'a|d'. Individual Ardu1 is represented by two extractions (r – root cementum, d – dentine).

The Kudruküla3 CCC hunter-gatherer sample was assigned to hg R1a5 as it was called with derived alleles in two R1a-M420 and one R1a5-YP1272 defining site while at 6 other R1a defining sites that were covered by data it showed the presence of an ancestral allele. Although the sample had low coverage, the hg R1a5 assignment of Kudruküla3 is further supported by the fact that it had 7 derived (versus one ancestral) alleles in upstream branches defining hg R while showing predominantly ancestral alleles at two other extant subclades of R. More specifically, we observed 11 ancestral calls and 1 derived call in R1b and 13 ancestral and no derived calls in R2 defining sites. These data support the assignment of Kudruküla3 into an early offshoot branch R1a5 of hg R1a. Our finding is further supported by the report of similar cases of R1a^43^ lineages which are derived at M459 but ancestral at M198 SNPs in two Eastern hunter-gatherer genomes, one from Karelia and the other from near Samara^2,4^. Altogether, these findings suggest that various subclades of R1a may have been common in hunter-gatherer populations of Eastern Europe and that just one of them, R1a-M417, was later amplified to high frequency by the Late Neolithic/Bronze Age expansion^2,25^. The fact that two Latvian hunter-gatherer Y chromosomes have been characterized as belonging to R1b-M269 clade^39^ suggests that both sub-clades of R1 were present in the Baltic area before the expansion of the CWC.

All four of the Estonian CWC individuals could be assigned to the R1a-Z645 sub-clade of hg R1a-M417 which together with N is one of the most common Y chromosome haplogroups in present-day Estonians (33%)^44^. Importantly, this R1a lineage is only distantly related to the R1a5 lineage we found in the CCC sample. The finding of high frequency of R1a-M417 in Estonian CWC samples is consistent with the observations made for other Corded Ware sites that, along with Late Bronze Age remains associated with Sintashta Culture, also show high frequency of hg R1a-M417^2,25^. However the remains associated with the Yamnaya Culture, which is considered to be ancestral to CWC, have been found with predominantly R1b-Z2105 and no R1a lineages^2,25^. On the other hand, no such clear differentiation was revealed when comparing the mtDNA haplogroup composition of the remains associated with the respective cultures^2,25^. The coalescent time for the R1a-Z645 clade, estimated from modern data at 5,400 yr BP (95% CI 4,950-6,000)^43^, predates the time when the CWC individuals carrying the R1a-Z645 lineages lived in Estonia (4,000–4,800 yr BP). The fact that all four of the CWC male individuals from two distinct sites in Estonia belonged to this recently expanded R1a branch, different from the one carried by CCC, suggests that admixture between CWC farmers and CCC hunter-gatherers may have been limited at least in the male lineages during the early stages of farming in Estonia. We acknowledge the limitations of our small sample size but, on the other hand, consider that this suggestion is supported by the similar findings made by Haak *et al.*^2^ and Lazaridis *et al.*^4^ for a wide range of CWC male samples.

### Estonian Neolithic among ancient and modern autosomal diversity

In order to reveal the place of the Estonian CCC and CWC aDNA samples on the canvas of the genome-wide genetic variation patterns in Europe, we first applied the Principal Component Analysis (PCA) approach integrating the newly-typed samples within the Human Origins dataset from Lazaridis *et al.* 2016^4^. We projected Estonian as well as other European and West Asian aDNA sequences^4^ to principal component axes constructed from the data of modern samples (Fig 2). Estonian CCC samples positioned between Scandinavian and Eastern hunter-gatherers (SHG and EHG) quite far from Western hunter-gatherers (WHG). Since the Mesolithic hunter-gatherer individuals from the Baltics have been shown to cluster with WHG^39,45^, this could point towards genetic input from the east. The Estonian CWC individuals on the other hand clustered closely together with a bulk of modern as well as Late Neolithic/Bronze Age (LNBA) populations from Europe.

**Fig 2.**
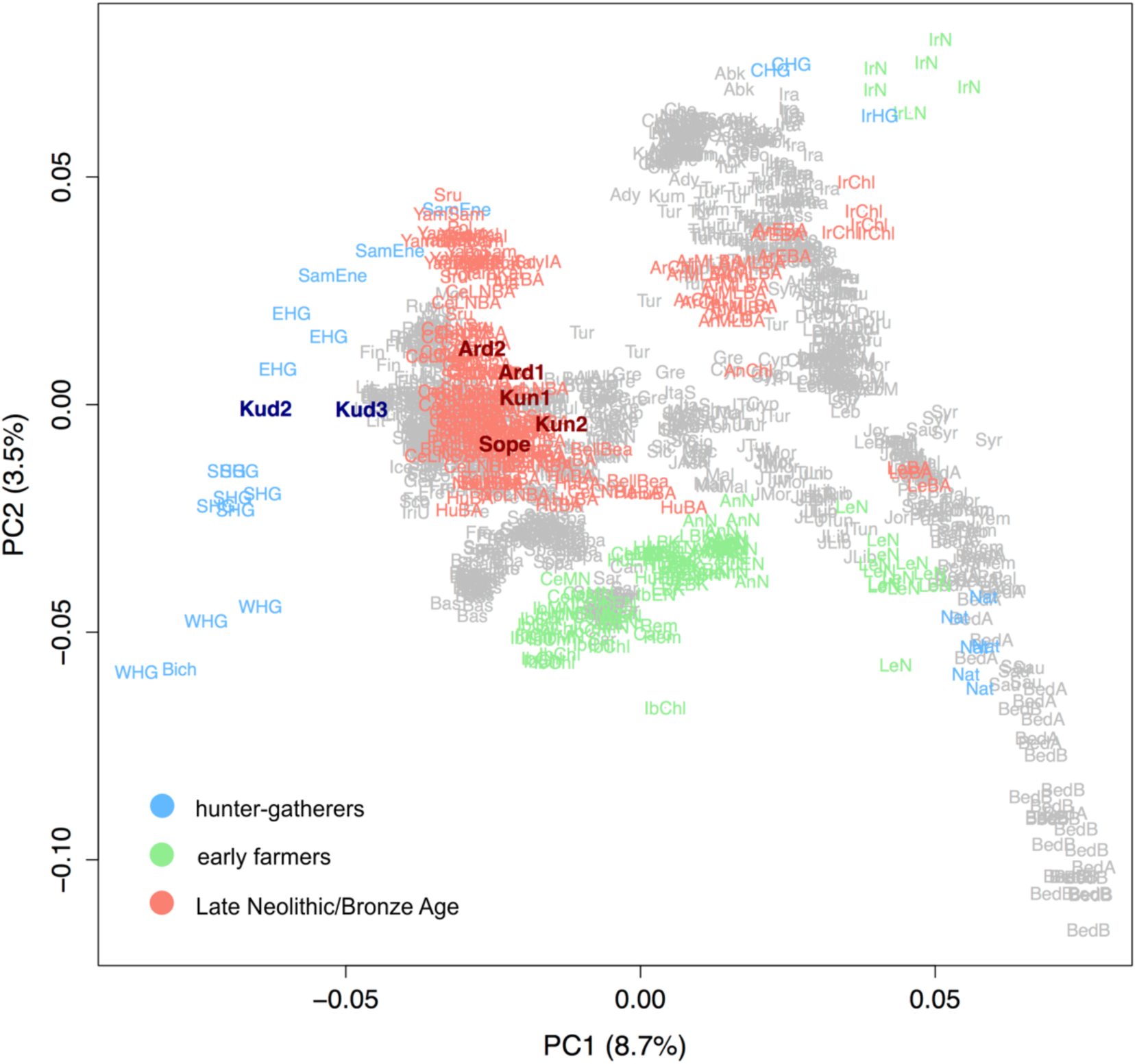
Principal component analysis of modern West Eurasians with ancient individuals (including 7 ancient Estonians) projected onto the first two components (PC1 and PC2).
CCC samples in dark blue, CWC samples in dark red. Kud2, 3 – Kudruküla; Ard1, 2 – Ardu; Kun1, 2 – Kunila.

ADMIXTURE^46^ analysis at K=14 (see Methods), where we projected aDNA samples on modern reference data^4^, revealed that Estonian CCC individuals were most similar to Eastern hunter-gatherers (Fig 3; S3 Figure) having predominantly a blue component with a minority of green and red. On the other hand, CWC showed a higher fraction of the green over the blue component and less of the red one. The blue component was ubiquitous to most European ancient and modern populations and predominant in WHG, EHG and SHG. The green component instead was maximized in Caucasus hunter-gatherers (CHG) and Iranians. Its southeast to northwest frequency decrease in Europe has been ascribed to the expansion of Yamnaya and Bronze Age cultures^2,25^. The red component detected in CCC, EHG and SHG individuals has its peak frequency in Eastern Siberian populations and is therefore likely to reflect ancient links between North Eurasian groups.

**Fig 3.**
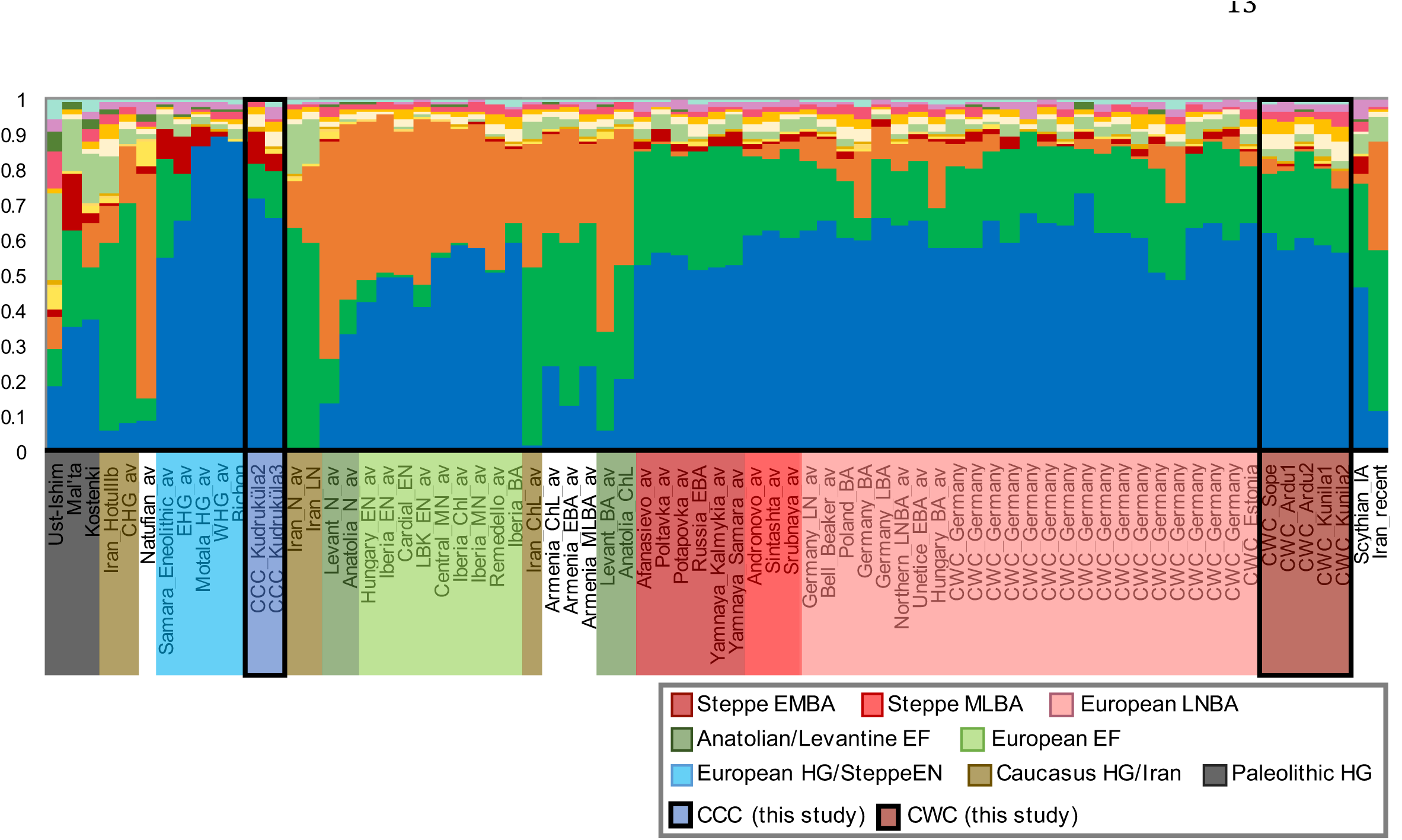
ADMIXTURE analysis results for aDNA samples at K14 with ancient individuals projected onto the modern genetic structure.The Y axis shows the proportions of the ancestral components.

Notably, the orange component characterizing early farming populations of Anatolia and Europe, which was also present in Middle/Late Bronze Age (MLBA) Steppe and European LNBA populations, was absent in all hunter-gatherer populations (including CCC) and Early/Middle Bronze Age (EMBA) Steppe populations (including Yamnaya). Two out of five Estonian CWC individuals had more than 3% of this component and these individuals also placed closer to early farmers on the PCA plot than the rest of Estonian CWC samples did (Fig 2). Another clear pattern that emerged while comparing EHG and CCC to LNBA populations was the lower prevalence of the red component in the latter. At this lower level there still seemed to be some structure to its distribution within LNBA. While it was present (1.9–5%, average 3.4%) in all Yamnaya samples it had an uneven distribution (0–3.5%, average 1.3%) in CWC samples. Interestingly, modern Estonians showed a bigger proportion of the blue component than CWC individuals. Comparing to CCC individuals, modern Estonians lack the red component. This, together with the absence of Y chromosome hg N in CCC and CWC, points to further influx and change of genetic material after the arrival of CWC.

Consistent with the ADMIXTURE results, both SHG and EHG were the closest to Estonian CCC samples also in outgroup f3 and Patterson’s D analyses^47^ followed by other Mesolithic hunter-gatherer and Steppe Eneolithic samples (Fig 4A and B, S4 Figure, S2 Table; S3 Table). By contrast, in the case of Estonian CWC, we do not see such a clear pattern of differentiation. Both Mesolithic hunter-gatherer and Late Neolithic to Bronze Age samples appear nearly equidistant from CWC, with EHG and SHG having marginally shorter distances (Fig 4A and B, S4 Figure, S2 Table; S3 Table). Interestingly, outgroup f3 in the form of f3(Yoruba; Yamnaya Samara, X) revealed that Estonian CCC, along with EHG and Steppe Eneolithic, shared more drift with Yamnaya Samara than did the LNBA populations of Europe (including Estonian CWC) (S5 Figure, S4 Table). This can be interpreted as sharing of EHG ancestry by Yamnaya and CCC populations on one hand and the Central European LNBA populations drawing additional ancestry from WHG and Anatolian farmers on the other. Notably, groups of Yamnaya samples from Samara and Kalmykia behaved similarly in these tests (S6 Figure, S5 Table).

**Fig4.**
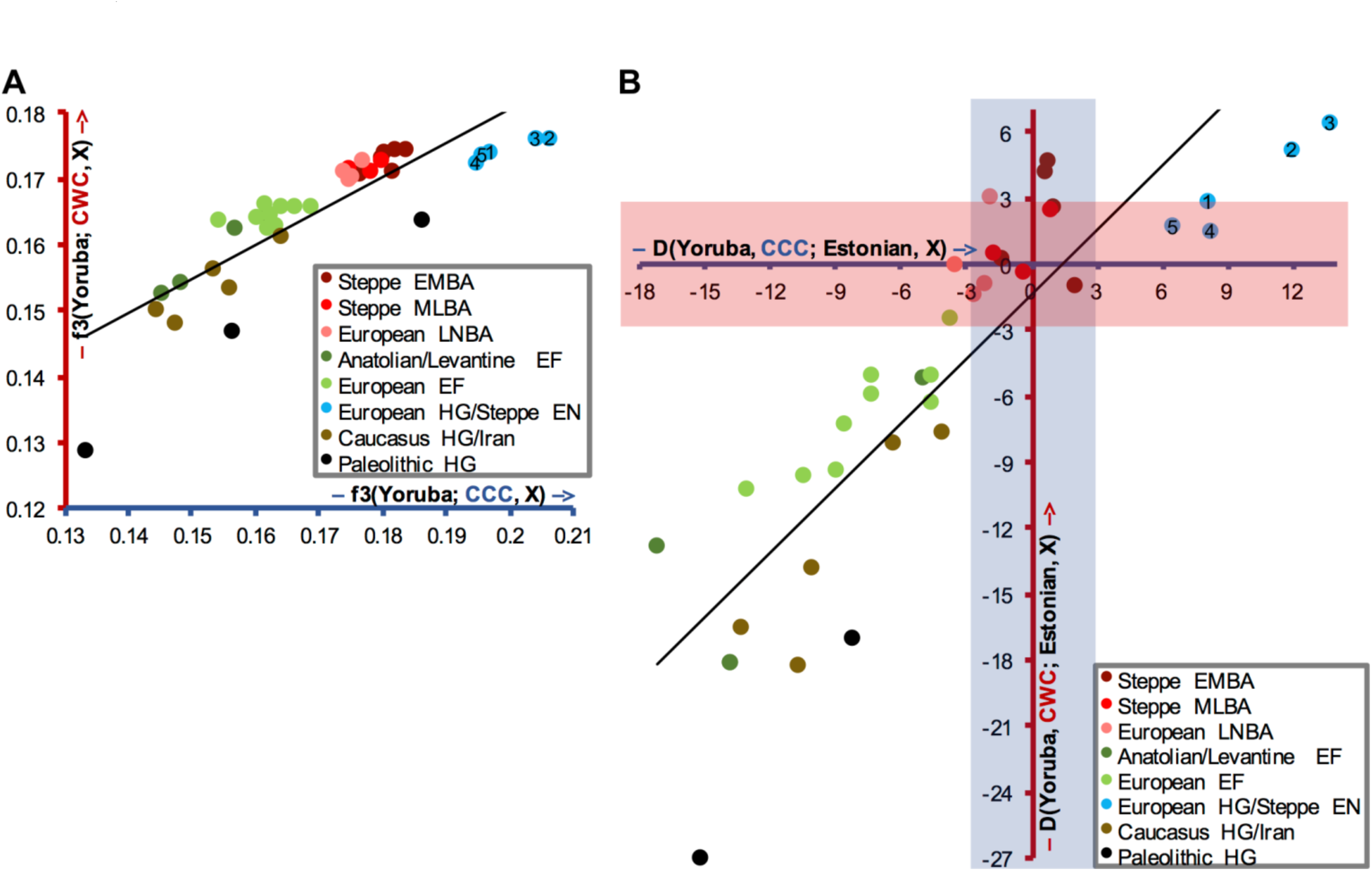
Outgroup f3 (A) or D statistics' (B) results of form f3(Yoruba; CCC/CWC, ancient) or D(Yoruba, CCC/CWC; Estonian, ancient) with Comb Ceramic Culture (CCC; blue axis) and Corded Ware Culture (CWC; red axis) plotted against each other. f3 values are used for A and Z scores for B. EMBA – Early/Middle Bronze Age; MLBA – Middle/Late Bronze Age; LNBA – Late Neolithic/Bronze Age; EF – early farmers; HG – hunter-gatherers; EN – Eneolithic. 1 – Steppe Eneolithic; 2 – Eastern HG; 3 – Scandinavian HG; 4 – Western HG; 5 – Switzerland HG.

When using modern populations from Europe, Caucasus, Near East and Siberia as test populations for calculating outgroup f3 statistics (S6 Table), we found that, among all present-day populations considered, both Estonian ancient groups were most similar to extant Lithuanians (Fig 5B). Also, modern populations speaking Uralic (including Finnic) languages of Europe and Siberia along with all other Siberians were more similar to Estonian CCC than to CWC (Fig 5). These results were confirmed by D statistic in the form D(Yoruba, CCC/CWC; Estonian, Y) (S7 Figure; S7 Table). This points to extant Siberians and European Uralic speakers having less Steppe ancestry than Estonian CWC did.

**Fig5.**
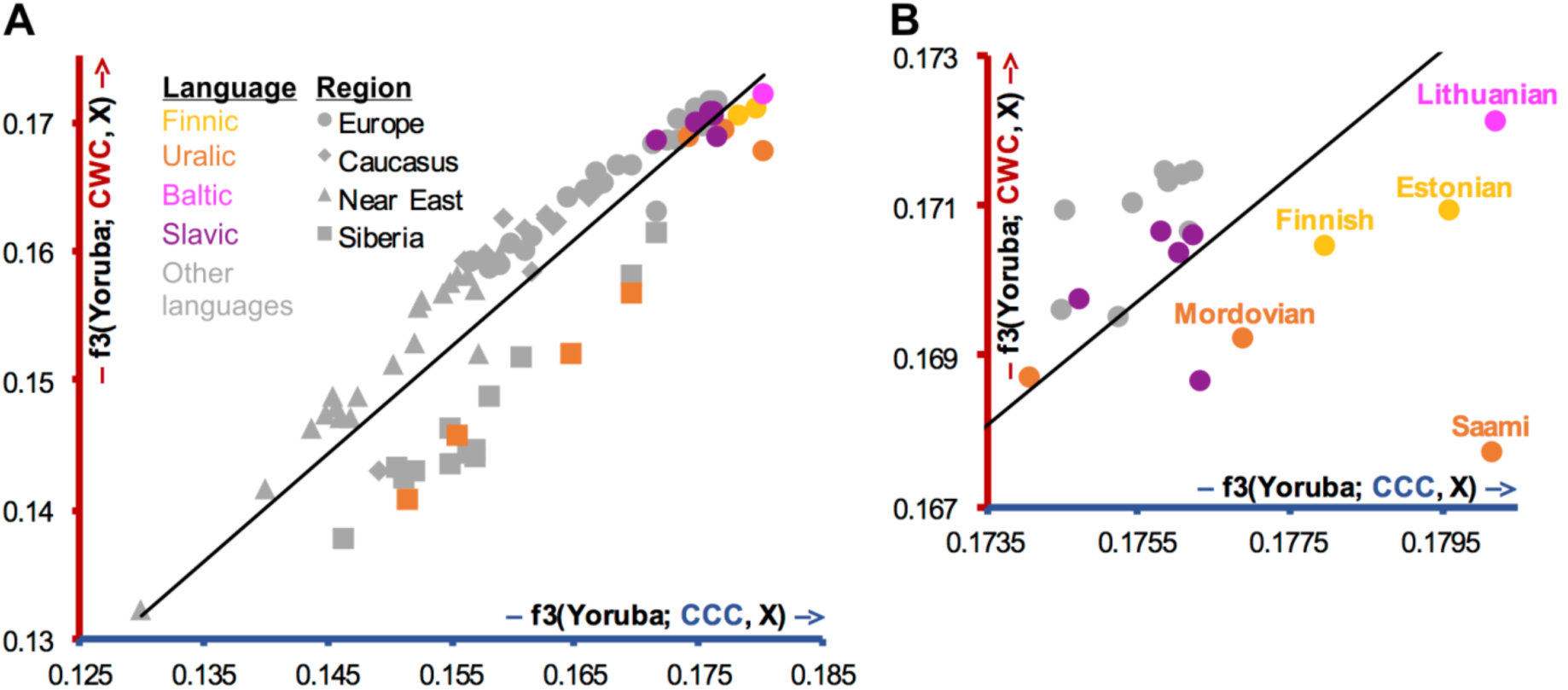
Outgroup f3 statistics' values of form f3(Yoruba; CCC/CWC, modern) with Comb Ceramic Culture (CCC; blue axis) and Corded Ware Culture (CWC; red axis) plotted against each other. A. Plot with all populations involved in the analyses. B. Top right corner of A. Colours signify languages and shapes signify regions.

Admixture f3 statistics with a subset of hunter-gatherer, early farmer and Late Neolithic/Bronze Age populations (S8 Table) revealed that Estonian CWC can be modelled through two types of admixture patterns. One combining Early Neolithic populations and EHG while in the other the mixing populations are CHG and any other hunter-gatherer group. It is interesting that Yamnaya, which in turn can be seen as a combination of CHG and EHG, is not directly needed for explaining the admixture pattern in Estonian CWC.

### Sex-specific ancestries of the Estonian Corded Ware Culture

Contrasting allele frequency patterns on autosomal (S9 Table) *vs* X chromosome(s) (S10 Table) in outgroup f3 analyses revealed that on X chromosome CWC is strikingly more similar to early farmer populations of both Anatolia/Levant and Europe. Accordingly, on the autosomes we see the opposite – most of the European hunter-gatherer/Steppe Eneolithic, EMBA Steppe and CHG/ancient Iranian populations tend to have higher f3 values (Fig 6). These results suggest that the gene flow associated with the spread of the CWC to Estonia may have been sex-specific, being biased toward Steppe ancestry in the male and early farmers on the female side. This finding is consistent with broader sex-specific patterns of admixture detected in European Bronze Age populations^48^.

**Fig 6.**
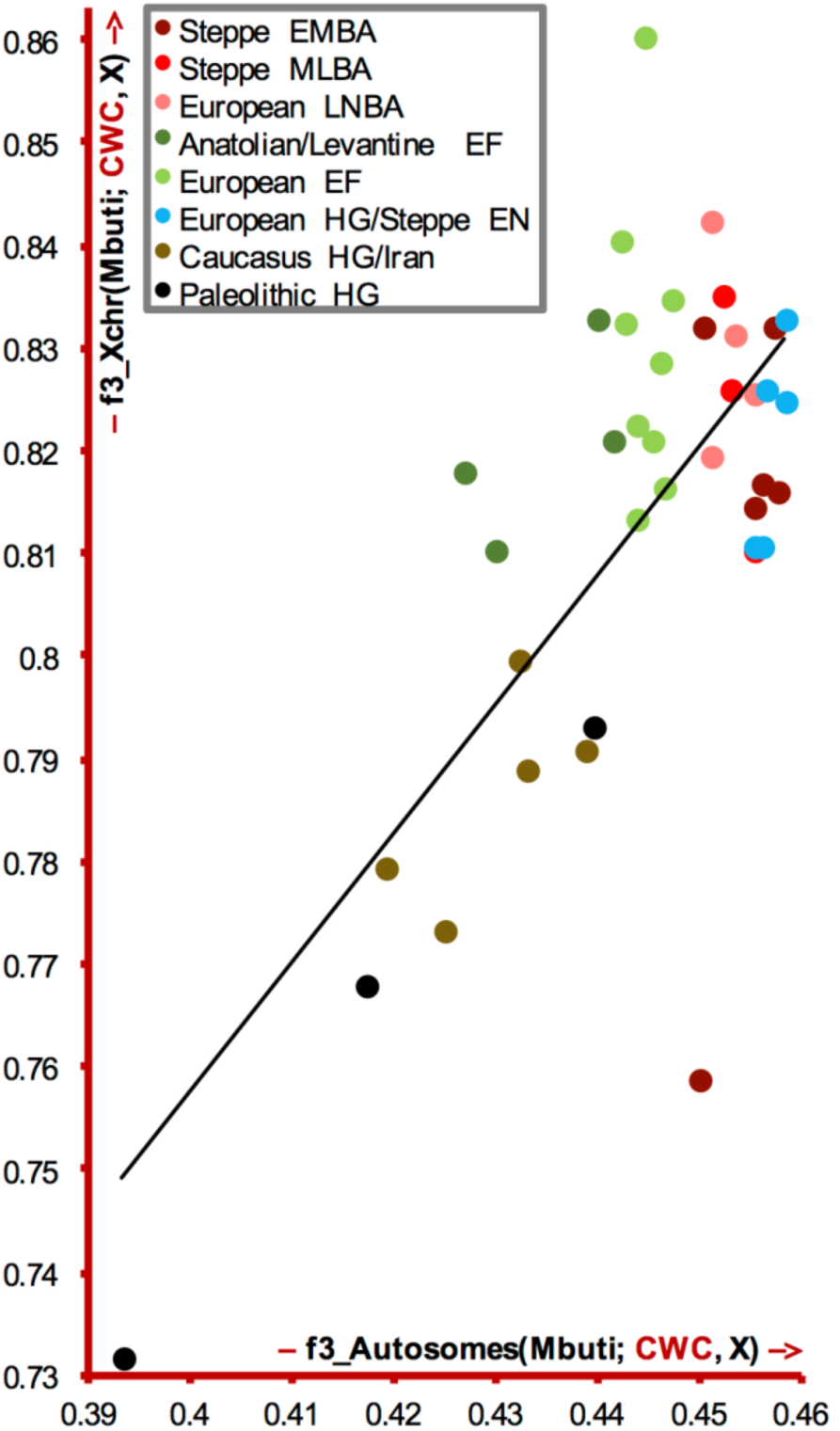
Outgroup f3 statistics' values of form f3_Xchr(Mbuti; CWC, ancient) and f3_Autosomes(Mbuti; CWC, ancient) plotted against each other. CWC – Corded Ware Culture. EMBA – Early/Middle Bronze Age; MLBA – Middle/Late Bronze Age; LNBA – Late Neolithic/Bronze Age; EF – early farmers; HG – hunter-gatherers; EN – Eneolithic.

## Conclusions

Our results support the hypothesis that individuals associated with the CCC hunter-gatherers in Estonia were genetically most similar to Eastern hunter-gatherers from Karelia, a region further east from Estonia. A recently published comparison of Latvian Mesolithic and CCC hunter-gatherer genomes supports this result^39^, while Mesolithic hunter-gatherers from Latvia and Lithuania appear genetically most similar to Western hunter-gatherers^39,45^. This implies a degree of genetic influx from the east with the arrival of CCC. Furthermore, the presence of a genetic component associated with Caucasus hunter-gatherers and later with people representing the Yamnaya Culture in Eastern hunter-gatherers and Estonian CCC individuals means that the expansion of the CWC cannot be seen as the sole means for the spread of this genetic component, at least in Eastern Europe. The transition to intensive farming and animal husbandry in Estonia, which took place a few thousand years after the farming transition in many other parts of Europe, was conveyed by the CWC individuals and involved an influx of new genetic material. These people carried a clear Steppe ancestry with some minor Anatolian contribution, most likely absorbed through female lineages during the population movements. Our results confirm a recently reported claim that farming did not arrive to the Baltic region as a consequence of step-wise migration of people from Anatolia^39^. However, our genetic data associated with the archaeological records demonstrate that the transition to farming in the Baltic region was not the result of only cultural transmission either but was instead the result of migration of farming people from the Steppe. What is more, since the genetic variation of contemporary Estonians cannot be fully explained by these events, further research will be needed to reveal the subsequent demographic events that brought Y chromosome hg N onto the shores of the Baltic Sea and changed the autosomal variation of the people living in this area.

## Materials and Methods

### Materials

The teeth used for DNA extraction were obtained with relevant institutional permissions from the Tallinn University Archaeological Research Collection and the Narva Museum.

DNA was extracted from 10 teeth, 5 of which belonged to Neolithic CWC individuals, 4 to Neolithic CCC individuals and 1 to a Mesolithic NC individual (Table 3). The carbon dated remains of CWC individuals are between 4,340 and 4,871 years old, of CCC individuals between 4,488 and 5,781 years old and the Mesolithic NC individual is between 6,361 and 6,236 years old. (Table 3). More detailed information about the archeological sites connected to each culture and the specific sites and burials of this study is given in S1 Appendix. Data from one of the CWC samples (Sope_r) was already included in the Bronze Age project The Rise^25^ under the sample name RISE00.

### DNA extraction, library preparation and DNA sequencing

All of the laboratory work was performed in dedicated ancient DNA laboratories at Centre for GeoGenetics, Natural History Museum, University of Copenhagen. For most of the samples, DNA was extracted, libraries prepared and DNA sequenced as in Allentoft *et al.,* 2015^25^. Since samples Kudruküla and Sope (1st extraction) were treated earlier, an older protocol with some differences was used while working with these samples. The main steps of the laboratory work and the differences between protocols are listed below.

### DNA extraction

The teeth were cleaned with a drill and cut in half horizontally. The cementum layer of the tooth roots was specifically targeted^49^ and for 3 teeth (1st extraction and sample Ardu1) DNA was also extracted from the dentine, giving 13 samples in total for DNA extraction.

To remove contaminants from the surface of tooth pieces/powder, a pre-digestion step was carried out by adding a custom digestion buffer^49^ and incubating the sample in a slow rotor for half an hour. After that the buffer was replaced with a clean one and the samples were left to digest over night. Undigested material was stored for a second DNA extraction if need be.

Silica beads and a custom binding buffer (as in Seguin-Orlando *et al.* 2014^50^ for 1st extraction, as in Allentoft *et al.* 2015^25^ for others) were added and the samples were kept in the dark in a rotor for approximately two hours to bind DNA to silica beads. Then the samples were washed with 80% ethanol and eluted using EB buffer (QIAquick PCR Purification Kit, QIAGEN).

### Library preparation

Sequencing libraries were built using NEBNext DNA Library Prep Master Mix Set 2 (E6070, New England Biolabs) and Illumina-specific adaptors^51^ following established protocols^51–53^. Two verification steps were implemented to make sure library preparation was successful – gel electrophoresis after PCR amplifications and a BioAnalyser (Agilent Technologies) run after purifying the libraries.

### DNA sequencing

DNA was sequenced using the Illumina HiSeq 2500 platform with the 100 bp single-end method. All samples were first sequenced together on one lane, one sample (Sope_r) became part of a Bronze Age project The Rise^25^ and was therefore sequenced many times, and 7 other samples from different individuals with endogenous DNA content over 1% were sequenced further on 8 lanes.

## Bioinformatics

### Mapping

Before mapping, the sequences of adaptors and indexes were cut from the ends of DNA sequences using Trimmomatic 0.32^54^ with the option ILLUMINACLIP. Sequences shorter than 30 bp were also removed with the option MINLEN to avoid random mapping of sequences from other species.

The sequences were mapped to reference sequence GRCh37 using Burrows-Wheeler Aligner (BWA)^55^ and command aln.

After mapping, the sequences were prepared for analyses by first converting them to SAM format with BWA command samse. Then the sequences were converted to BAM format, sequences that mapped to the reference sequence were sorted out, and PCR duplicates were removed, all of which was done with samtools 0.1.19^56^.

The average endogenous DNA content (proportion of reads mapping to the human genome) for hunter-gatherers was 2% but for farmers 36.5% (Table 4). Eight samples with the highest endogenous DNA content were sequenced further (* in Table 4).

### aDNA authentication

As a result of degrading over time aDNA can be distinguished from modern DNA by certain characteristics: short fragments with long single-stranded overhangs and a high frequency of C=>T substitutions at the 5' ends of sequences due to cytosine deamination. The program mapDamage2.0^57^ was used to estimate the frequency of 5' C=>T transitions, the fraction of bases located in single-stranded overhangs (λ) and the C=>T substitution rate in those overhangs (*δ*s).

mtDNA contamination was estimated using the method from Fu *et al.* 2013^7^.This included calling an mtDNA consensus sequence based on reads with mapping quality at least 30 and positions with at least 5x coverage, aligning the consensus with 311 other human mtDNA sequences from Fu *et al.* 2013^7^, mapping the original mtDNA reads to the consensus sequence and running contamMix 1.0-10 with the reads mapping to the consensus and the 312 aligned mtDNA sequences while trimming 7 bases from the ends of reads with the option trimBases.

For the male individuals we also estimated contamination based on X chromosome using the contamination estimation method first described in Rasmussen *et al.* 2011^58^ and incorporated in the ANGSD software^59^.

The samples showed C=>T substitutions at the 5' ends 8–24% (Table 4), λ 31–45% and δs 18–66%. mtDNA contamination could be estimated for 10 out of the 13 samples and it's point estimate ranged from 0.1% to 2.5% (Table 4). X chromosome contamination could be estimated for 3 of the 7 male individuals and was between 0.9% and 1.9% (Table 4).

### Calculating coverage and determining genetic sex

BEDTools 2.19.0 genomecov^60^ was used to determine the coverage of each nucleotide in the reference sequence. Mitochondrial and full genome coverages were calculated as the average of the nucleotide coverages of each genome.

Genetic sex was calculated as in Skoglund *et al.* 2013^61^, estimating the fraction of reads mapping to Y chromosome out of all reads mapping to either X or Y chromosome.

The mtDNA coverage for all of the samples was between 0.07 and 130 and the genomic coverage ranged from 0.00006 to 2.13 (Table 4). Genetic sexing confirmed morfological sex estimates (where available) and provided additional information about the sex of the individuals involved in the study. The sex of one of the samples, Naakamäe could not be estimated due to low coverage. Apart from the sample Naakamäe, the study involves 3 females and 6 males (Table 4).

### Variant calling

Variants were called with the ANGSD software^59^ command doHaploCall, sampling a random base for the positions that are present in the Lazaridis *et al.* 2016^4^ aDNA dataset. Variants were called for each sample separately and also for merged samples from the same individual (merged with samtools merge^56^).

### Determining mtDNAhaplogroups

mtDNA haplogroups were determined by submitting mtDNA VCF files to HaploGrep2^62,63^. Later the results were checked visually by aligning mapped sequences to reference sequence rCRS^64^ with samtools 0.1.19^56^ command tview and comparing the variants with the previously suggested lineage in PhyloTree^63^.

### Y chromosome variant calling and haplogrouping

Y chromosome variants were called from the bam files of the six male samples that had higher than 0.1x genome coverage, five from CWC and one from CCC group, using ANGSD^59^. The resulting VCF files were filtered for regions of a total length of 8.8 Mbp sequence that uniquely maps to human Y chromosome when using short read sequencing technology^43^. Variants called within this 8.8 Mbp region were further filtered for 42,385 haplogroup informative markers^43^ using BEDTools 2.19.0 intersect option. Haplogroup assignments of each individual sample were made by determining the haplogroup with the highest proportion of markers called in derived state in the given sample.

### Principal component analysis

To prepare for principal component analysis (PCA), the per-individual data of this study was converted to BED format using PLINK 1.90 (http://pngu.mgh.harvard.edu/purcell/plink/>)^65^. Data from 991 individuals from 67 populations of Europe, Caucasus and Near East from the Human Origins dataset^4^ (S11 Table) was used as the modern DNA background. 276 samples from 25 populations from Lazaridis *et al.* 2016^4^ were used as an aDNA comparison dataset (S11 Table), leaving out the recent Iranian and Sope (RISE00) individual. The 3 datasets were merged and only the positions from the Human Origins dataset that were used in Lazaridis *et al.* 2016^4^ were kept (587,469 SNPs). Due to a low number of SNPs, 3 of the ancient Estonian individuals were removed from further autosomal analyses, leaving 2 CCC (Kudruküla2 and 3) and 5 CWC individuals (Ardu1 and 2, Kunila1 and 2, Sope) from Estonia to be used in autosomal analyses. The data was converted to EIGENSTRAT format using the program convertf from the EIGENSOFT 5.0.2 package^66^. PCA was performed with the program smartpca from the same package, projecting ancient samples onto the components constructed based on the modern dataset using the option lsqproject.

To see how much the subset of SNPs present in a given aDNA sample influences its placement on the background of modern samples on the PCA plot, a test was run using different subsets of SNPs of a modern sample. The exact SNPs and ten sets of the same number of random SNPs as are present in the two lowest covered samples (7,773 and 33,488 SNPs) and the best covered sample (497,557 SNPs) of this study were used. It was evident that all of these new samples placed around the original modern sample and in the same region but while the best covered Estonian aDNA sample's SNPs allowed the new samples to fall almost exactly on top of the original sample, the worst covered samples' SNPs did not allow for interpretation of their exact placement (S8 Figure).

### Admixture

2273 modern individuals from 226 populations from the Human Origins dataset^4^, 278 ancient samples from Lazaridis *et al.* 2016^4^ (including the recent Iranian and Sope individual), and 7 Estonian samples from this study were included into Admixture^46^ analysis. The analysis was carried out using ADMIXTURE 1.3^46^ with the P option, projecting the ancient samples into the genetic structure calculated on the modern dataset. The dataset of modern samples was pruned to decrease linkage disequilibrium using the option indep-pairwise with parameters 1000 250 0.4 in PLINK 1.90 (http://pngu.mgh.harvard.edu/purcell/plink/)^65^. This resulted in a set of 297 478 SNPs. We ran Admixture on this set using K=3 to K=18 in 100 replicates. This enabled us to assess convergence of the different models. K=14 was the model with the largest number of inferred genetic clusters for which >10% of the runs that reached the highest Log Likelihood values yielded very similar results. We use this as a proxy to assume that the global Likelihood maximum for this particular model was indeed reached. We then used the inferred genetic cluster proportions and allele frequencies of the best run at K=14 and used these to run Admixture to project the aDNA samples on the inferred clusters. We present only these aDNA samples for which the intersection with the LD pruned modern samples dataset yielded data for more than 10 000 SNPs. The resulting membership proportions to K genetic clusters are sometimes called “ancestry components” which can lead to over-interpretation of the results. The clustering itself is, however, an objective description of genetic structure and as such a valuable tool in population comparisons.

### Outgroup f3 statistics

Data from extant populations of Europe, Siberia and Africa from the Lazaridis *et al.* 2016 Human Origins dataset^4^ was added to the dataset used for PCA, making the f3 modern dataset comprise of 1309 individuals from 89 populations (S11 Table). 255 ancient individuals from 23 populations were used in the analyses, including 3 Paleolithic hunter-gatherers (S11 Table). Heterozygous positions were converted to homozygous by randomly choosing one of the alleles at each position to enable comparison between pseudo-haploid ancient samples and diploid modern samples. The data was converted to EIGENSTRAT format using the program convertf from the EIGENSOFT 5.0.2 package^66^. Outgroup f3 statistics of the form f3(Yoruba; CCC/CWC, modern/ancient) were computed using ADMIXTOOLS 1.1^47^.

To allow for X chromosome *versus* autosomes comparison, outgroup f3 statistics of the form f3(Mbuti; CCC/CWC, ancient) were computed with Mbuti from Panel C of the Simons Genome Diversity Project^67^ as the outgroup. The analyses were run both using the same autosomal SNPs as before, and also all X chromosome positions available in the ancient dataset (49,075 SNPs). Since all children inherit half of their autosomal material from their father but only female children inherit their X chromosome from their father then in this comparison X chromosome data gives more information about the female and autosomal data about the male ancestors of a population.

### D statistics

D statistics of the form D(Yoruba, CCC/CWC; Estonia, modern/ancient) were calculated on the same dataset as outgroup f3 statistics (S11 Table). The program POPSTATS^68^ with the option informative was used.

### Admixture f3 statistics

To compute admixture f3 statistics, a subset of ancient populations was chosen – Natufian, Caucasus, Eastern, Motala and Western hunter-gatherer, Steppe Eneolithic, Anatolia Neolithic, LBK Early Neolithic, Yamnaya Kalmykia, Yamnaya Samara, Central Late Neolithic/Bronze Age, and Estonian CCC and CWC. f3 statistics were calculated using the SNPs of the ancient dataset (1,047,341 SNPs) for all combinations this time using the non-homozygous data and ADMIXTOOLS 1.1^47^ with the parameter inbreed: YES.

Due to a low number of informative positions there were no significant results when Estonian CCC was involved in the analysis (S8 Table).

## Acknowledgements

This work was funded by research projects of the Estonian Research Council IUT24-1 (T.K., E.M., M.M., M R.,La.S, K.T.), IUT20-7 (A.K.) and PUT1217 (K.T., La.S.,Le.S., L.V.); EU European Regional Development Fund Centre of Excellence in Genomics and Translational Medicine TK142 and Estonian Centre for Genomics 2014-2020.4.01.16-0125; ERC Starting Investigator grant FP7-261213 (T.K.) and Arheograator Ltd. (L.V., A.K.)

## Author contributions

Le.S., E.M., T.K., A.K. and M.M. conceived the study. L.V. and A.K. assembled the skeletal samples and performed osteological analyses. Le.S., J.S., M.A. and E.W. did aDNA extraction and DNA sequencing. Le.S., C.L.S., La.S., L.P., M.R., E.M., T.K. and M.M. analysed the data. Le.S., L.P., K.T., L.V., E.M., A.K., T.K. and M.M. wrote the manuscript with input from the rest of the authors.

## Competing financial interests

The authors declare no competing financial interests.

### Materials and Correspondence

The DNA sequences are available through the data depository of the EBC http://www.ebc.ee/free_data following the Fort Lauderdale principle.

Correspondence is welcome to LehtiSaag and Mait Metspalu

